# The role of preterm birth and postnatal stress in neonatal structural brain development

**DOI:** 10.1101/2022.01.28.478158

**Authors:** Femke Lammertink, Manon J.N.L. Benders, Erno J. Hermans, Maria L. Tataranno, Jeroen Dudink, Christiaan H. Vinkers, Martijn P. van den Heuvel

## Abstract

Preterm birth disrupts the emerging foundations of the brain’s architecture, and the continuum of early-life stress-provoked alterations reaches from a healthy adaptation with resilience to severe vulnerability and maladjustment with psychopathology. The current study examined how structural brain development is affected by a stressful extra-uterine environment and whether changes in topological architecture at term-equivalent age could explain the increased vulnerability for behavioral symptoms during early childhood. Longitudinal changes in structural brain connectivity were quantified using diffusion-weighted imaging (DWI) and tractography in preterm born infants (gestational age <28 weeks), imaged at 30 and/or 40 weeks of gestation (*N*=145, 43.5% female). A global index of postnatal stress was based on invasive procedures during hospitalization (e.g., heel lance). Infants were classified as vulnerable and resilient based on having more or less internalizing symptoms at 2-5 years of age (*n*=71). Findings were replicated in an independent validation sample (*N*=123, 39.8% female, *n*=91 with follow-up). Higher stress levels impaired structural connectivity growth in the amygdala, insula, hippocampus, and posterior cingulate cortex. The hippocampus, amygdala, and subthalamic nucleus showed lower global connectivity in vulnerable relative to resilient individuals. The distinct characteristics of the resilient brain allowed for a good predictive accuracy of group membership using local network measures (80%, *p*<10^−5^, κ=0.61). These findings emphasize the detrimental impact of postnatal stress and, more importantly, the relative plasticity of the preterm brain. Resilience following postnatal stress appertains to a potential compensatory or innate ability to propagate global information flow.

**Significance Statement:** The underdeveloped preterm brain is exposed to various external stimuli following birth. Although the importance of early adversity has been widely recognized, the essential understanding of the effects of early chronic stress on neonatal brain networks as well as the remarkable degree of resilience is not well understood. We aim to provide an increased understanding of the impact of postnatal stress on brain development between 30 and 40 weeks of gestation and describe the topological architecture of a resilient brain. We observed global alteration in neonatal brain networks following postnatal stress and identified key contributive regions conferring resilience to the development of future internalizing symptoms.

## Introduction

During critical periods of brain development, the extra-uterine environment impacts the maturation of the structural brain and behavioral functions. Preterm birth has long-lasting adverse effects on brain development and increases the risk for psychiatric symptoms later in life (Eikenes et al., 2011; Fischi-Gómez et al., 2015; Loe et al., 2013; Spittle et al., 2009). The development of the preterm brain is contingent on several (clinical) factors, and emerging data suggest that postnatal stressors such as the number of invasive procedures also play a role (Chau et al., 2019; Doesburg et al., 2013; Ranger et al., 2015; Ranger & Grunau, 2013). A paucity of longitudinal studies has explored the complex interaction between postnatal stress, brain development, and behavioral functions following preterm birth. We thus examined the impact of extra-uterine postnatal stress on brain development and how alterations in brain network architecture influences vulnerability for behavioral symptoms during early childhood (2-5 years).

Early-life adversities may alter trajectories of brain maturation during a critical period of development. Cross-sectional studies investigating the effects of preterm birth on brain structure and function have shown lower white matter integrity in association tracts (forceps minor, forceps major, inferior frontal-occipital fasciculus/inferior longitudinal fasciculus, superior longitudinal fasciculus, and uncinate fasciculus); and projection fibers (e.g., thalamic radiation, corticospinal tract; Duerden et al., 2018; Menegaux et al., 2017; Vollmer et al., 2017; Zwicker et al., 2013). Preterm birth is further related to an upregulation of functional connectivity between stress-related and stress-vulnerable regions, such as the temporal cortex, thalamus, anterior cingulate gyrus, hippocampus, and amygdala (De Asis-Cruz et al., 2020; Johns et al., 2019; Papini et al., 2016). More recently, advances in graph theory enabled researchers to reveal meaningful information about the topological architecture of the neonatal brain. Studies showed, for instance, that the fundamental community structural properties (i.e., groups of densely connected regions reflecting subsystems or “building blocks” of a network) of a preterm born infant seem to be similar to typically developing fetuses and neonates (Song et al., 2017; Turk et al., 2019); initial connectomic studies also highlight a more segregated and less integrated network organization in preterm-born infants (Ball, Boardman, et al., 2013; Ball, Srinivasan, et al., 2013; Groppo et al., 2014; Sa de Almeida et al., 2021) and children (de Kieviet et al., 2021; Fischi-Gomez et al., 2016), indicating differences in connectomic composition. These neonatal alterations in brain connectivity architecture may play a significant role in developing future psychopathology (Gilchrist et al., 2021; Kaufmann et al., 2017; Van Essen & Barch, 2015). Indeed, altered brain connectivity is implicated in a wide range of major psychiatric conditions, from ADHD and anxiety to Major Depressive Disorder (Suo et al., 2017; Tozzi et al., 2021; Wang et al., 2021).

In this study, we examined the influence of stress on the development of premature brain connectivity and, second, whether alterations in macroscale network architecture at term-equivalent age may be predictive of vulnerability for anxiety-related symptoms during early childhood (2-5 years of age). We examined diffusion imaging and tractography from preterm infants, combined with data on postnatal stress related to their hospitalization. We aim to identify specific differences in resilient and vulnerable infants that may enable resilient individuals to maintain relative mental wellbeing during early childhood.

## Materials and Methods

### Subjects

Infants were included when they were scanned between 28-32 and/or 39-42 post-menstrual age. Data collection was part of standard clinical care, with permission obtained to use this data for clinical research from the medical ethical review committee of the University Medical Center Utrecht (METC Utrecht). Preterm infants with chromosomal and/or congenital anomalies were excluded. Details and demographics of the main and validation datasets are outlined in Table 1.

**Table 1.** Sample demographic and neonatal clinical details of participants (N = 268)

| **Demographics** | **Main**<br>**Protocol A**<br>**(N = 145)** | **Validation**<br>**Protocol B**<br>**(N = 123)** |
| --- | --- | --- |
| Age at birth, mean ± SD, weeks | 26.53±1.01 | 26.54±1.00 |
| Age at scan, mean ± SD, weeks | 31.00±0.84<br>41.27±0.68 | 30.71±0.84<br>41.33±1.01 |
| 30 week MRI, n | 76 | 55 |
| 40 weeks MRI, n | 128 | 110 |
| serial MRI, n | 59 | 42 |
| Gender, female/male, n | 63/82 | 49/74 |
| Birthweight z-score<sup>a</sup>, mean ± SD, grams | -0.61±1.41 | -0.52±1.44 |
| Postnatal stress<sup>b</sup>, median [range] | -0.69 (-3.26-5.14) | -1.26(-1.59-4.80) |
| Days of morphine, mean ± SD | 3.72±7.11 | 3.17±5.18 |
| Prenatal corticosteroids [yes/no] | 128/17 | 115/8 |
| Postnatal corticosteroids [yes/no] | 39/106 | 40/83 |
| Intraventricular hemorrhaging [yes/no] | 44/101 | 37/85 |
| Necrotizing enterocolitis, n | 21 | 10 |
| Retinopathy of prematurity, n | 53 | 40 |

*nb. Protocol A refers to 45 diffusion directions, Protocol B refers to 32 diffusion directions.*

#### Main dataset

Data of *N*=145 preterm infants born infants clinically diagnosed as ‘extremely preterm’ with a gestational age <28 weeks were included in our study, admitted to the Neonatal Intensive Care Unit (NICU) between 2013 and 2019 at the Wilhelmina Children’s Hospital Utrecht, The Netherlands. Infants were scanned using a 45 directions diffusion protocol.

#### Validation dataset

A replication sample containing *N*=123 preterm infants born infants with a gestational age <28 weeks was included to assess the robustness of our results. Infants were admitted to the NICU between 2008 and 2013 and were scanned using a 32 directions diffusion protocol.

### Magnetic Resonance Imaging

MRI data included the examination of 3T structural anatomical T2-weighted imaging and diffusion-tensor imaging (main dataset: dMRI, n=45 directions; validation dataset, n=32 directions) (3T Achieva MR scanner). Images were obtained as part of a 35-minute scanning session.

T2 data were acquired using a Turbo Spin Echo (TSE) sequence, using parameters: TR=6112ms, TE=120ms, voxel resolution in millimeters 0.53×0.64×2 for 30 weeks and TR=4851ms, TE=150ms, voxel resolution in millimeters 0.78×0.89×1.2 for 40 weeks. dMRI data were acquired at 2 mm isotropic resolution and SENSE factor of 2 in 2 shells; 45 non-collinear directions for the main dataset, with a b-value of 800 s/mm^2^ and one non-diffusion weighted image (b=0) with TR 6500 ms and TE 80 ms; and 32 non-collinear directions for the validation dataset, with a b-value of 800 s/mm^2^ and one non-diffusion weighted image (b=0) with TR 5685 ms and TE 70 ms.

Infants were immobilized by wrapping them into a vacuum cushion. MiniMuffs (Natus Europe, Münich, Germany) and earmuffs (EM’s kids Everton Park, Australia) were used to reduce noise and the infant’s propensity to move during image acquisition. Before scanning, preterm born infants scanned at 30 weeks were either sedated with 30 mg/kg oral chloral hydrate or not sedated at all, whereas infants scanned at 40 weeks were all sedated with 50 to 60 mg/kg oral chloral hydrate. Scanning was halted if the infant woke up, and attempts were made to re-settle the infant without taking them out of the patient immobilization system. A neonatologist or physician assistant was present at all times during the examination.

### Data processing

#### Structural images

Volumetric tissue segmentation of grey and white matter, and labeling of subcortical and cortical areas, was performed on the T2 image (voxel resolution in millimeters 0.53 × 0.64 × 2 for 30 weeks and 0.78 × 0.89 × 1.2 for 40 weeks) using the structural pipeline from the developmental human connectome project (dHCP; http://www.developingconnectome.org/). The dHCP pipeline utilizes an “Expectation-Maximization” scheme that combines structure priors and an intensity model of the images (Makropoulos et al., 2018). A total of 47 (sub-)cortical grey matter labels were automatically generated during segmentation (see Figure 1).

**Figure 1.**
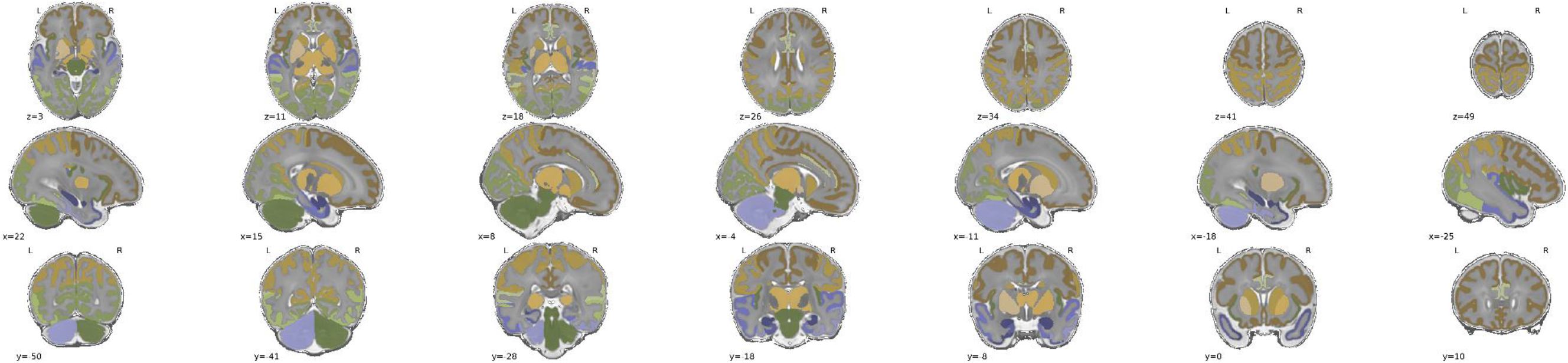
A total of 47 grey matter regions are segmented by the structural pipeline of the developmental Human Connectome Project (dHCP). HPL; hippocampus left, HPR; hippocampus right, AML; amygdala left, AMR; amygdala right, ATLML; anterior temporal lobe medial part left, ATLMR; anterior temporal lobe medial part right, ATLLL; anterior temporal lobe lateral part left; ATLLR; anterior temporal lobe lateral part right, GPAL; gyri parahippocampalis et ambiens anterior part left, GPAR; gyri parahippocampalis et ambiens anterior part right, STGL; superior temporal gyrus middle part left, STGR; superior temporal gyrus middle part right, MITGAL; medial and inferior temporal gyri anterior part left, MITGAR; medial and inferior temporal gyri anterior part right, LOGAL; lateral occipitotemporal gyrus/anterior fusiform left, LOGAR; lateral occipitotemporal gyrus/anterior fusiform right, CBL; cerebellum left, CBR; cerebellum right, BRS; brainstem, OLL; occipital lobe left, OLR; occipital lobe right, GPPL; gyri parahippocampalis et ambiens posterior part left; GPPR; gyri parahippocampalis et ambiens posterior right, LOGPL; lateral occipitotermporal gyrus/posterior fusiform part left, LOGPR; lateral occipitotermporal gyrus/posterior fusiform part right, MITGPL; medial and inferior temporal gyri posterior part left, MITGPR; medial and inferior temporal gyri posterior part right, STGPL; superior temporal gyrus posterior part left, STGPR; superior temporal gyrus posterior part right, CGAL; cingulate gyrus anterior part left, CGAR; cingulate gyrus anterior part right, FLL; frontal lobe left, FLR; frontal lobe right, PLL; parietal lobe left, PLR; parietal lobe right, CNL; caudate nucleus left, CNR; caudate nucleus right, THL; thalamus left, THR; thalamus right, SNL; subthalamic nucleus left, SNR; subthalamic nucleus right, LNL; lentiform nucleus left, LNR; lentiform nucleus right.

#### DWI tractography

Diffusion-weighted images were corrected for eddy current distortions, motion-induced signal drop-out, and head motion using a non-parametric approach using FSL (FSL EDDY) (Andersson & Stamatios, 2016). The b0 image (voxel-size 2×2×2 for the main dataset, voxel-size 1.41×1.41×2.00 for validation dataset, b = 0 s/mm^2^) was registered to the T2-weighted image for anatomical alignment of the DWI images using FLIRT with a boundary-based-registration (BBR) cost function (Greve & Fischl, 2009). The linear transformation matrix was combined with a non-linear warp registration using FSL FNIRT (Andersson et al., 2007) to map the diffusion space to an age-matched template. A single tensor model was used to estimate the main diffusion direction in each voxel (Basser et al., 1994) based on the 45 diffusion-weighted images (b = 800 s/mm^2^; 32 directions for the validation dataset). An FA and MD whole-brain map was created based on the fitted tensors. White matter pathways were reconstructed using FACT (fiber assignment by continuous tracking [Mori & Van Zijl, 2002]). Tractography involved starting eight streamline seeds in each white matter voxel, with fiber tracking, continued along the main diffusion direction of each voxel until a streamline showed high curvature (>65°), exited the brain mask, and/or when a streamline entered a voxel with low FA (<0.05). The mean FA value of a streamline was computed as the weighted average FA value, including all voxels that a streamline passed. Individual brain networks consisting of 47 grey matter regions and their interconnecting pathways were created by combining the subcortical and cortical segmentation map with all reconstructed white matter tractography streamlines, mapping for all combinations of regions their interconnecting streamlines, with the weight of each region-to-region connection taken as the non-zero mean FA of the selected streamlines. Connections with a low connectivity strength (lowest 5%) were taken as potential false-positive reconstructions and set to 0. A group-based threshold was applied, retaining connections present in at least 50% of the participants, balancing the number of false-positive and false-negative structural connections (de Reus & van den Heuvel, 2013). Results were validated using different levels of group-based consensus thresholds (50-90%, steps of 5%).

Three summary measures were used to detect outliers among connectivity matrices, namely the presence of odd connections, the absence of common connections, and the average fractional anisotropy. We calculated the interquartile range (IQR) for each group separately by subtracting the 25th percentile from the 75th percentile (i.e., IQR = Q3-Q1). Participants with a score below Q1-2×IQR or above Q2+2×IQR for any of the three measures were considered outliers. This quantification led to the removal of 7 outliers at 30 weeks of gestation and 19 outliers at 40 weeks of gestation.

### Behavioral measures

#### Postnatal stress

Data on invasive and stressful procedures were automatically extracted from the digital medical system. A global index of NICU-related stress was computed using a Principal Component Analysis on six parameters: skin-breaking procedures (i.e., heel lance, arterial and venous punctures, peripheral venous line insertion), total days of invasive mechanical ventilation, and suctioning of the nose and mouth. Each row (i.e., subject) was weighted on the total days of NICU admission. The extracted component explained 72.5% of the variance, with factor loading ranging from 0.74 to 0.91. This approach avoids the confounding effects of multicollinearity and continuously measures global NICU-related stress in further analyses.

#### Residualized approach to postnatal stress

All participants were invited for standard clinical follow-up at 2.5 and/or 5.75 years of age. Both the main and validation dataset had follow-up data, resulting in a total of 162 infants with both an MRI a term-equivalent age and data on behavioral symptoms (see Table 2 for an overview). During the clinical follow-up, parents reported on the level of internalizing symptoms of their child, such as depression and anxiety, using the Child Behaviour Checklist (CBCL; Achenbach & Rescorla, 2001). The CBCL is a parent-report questionnaire used to assess the frequency of dysfunctional behavior exhibited by the child in the past six months. Caregivers rate their children’s behavior by answering questions about their child on a 3-point scale (0-2), zero being “not true” one being “somewhat or sometimes true”, and two being “very true or often true”. If children did not have a behavioral symptom assessment at 5.75 years of age, we used the 2.5 years assessment (moderate correlation between the two-time points; *r*= 0.45, *p* < 0.001, see Figure 2-A). The follow-up also included other assessments not part of the current study, such as motor development and intelligence.

**Table 2.** Sample demographic and neonatal clinical details of resilient and vulnerable infants

| **Demographics** | **Main dataset** |  |  | **Validation dataset** |  |  |
| --- | --- | --- | --- | --- | --- | --- |
|  | **Resilient**<br>**(N = 41)** | **Vulnerable**<br>**(N = 30)** | **p-value** | **Resilient**<br>**(N = 42)** | **Vulnerable**<br>**(N = 49)** | **p-value** |
| Age at birth, mean ± SD, weeks | 26.63±1.00 | 26.54±0.92 | ns | 26.47±1.00 | 25.57±0.92 | ns |
| Age at scan, mean ± SD, weeks | 41.17±0.78 | 41.22±0.46 | ns | 41.14±0.48 | 41.46±1.36 | ns |
| Gender, female/male, n | 12/29 | 16/14 | < 0.05 | 16/26 | 14/35 | ns |
| Birthweight z-score<sup>a</sup>, mean ± SD, grams | -0.44±1.35 | -0.82±1.47 | ns | -0.61±1.31 | -0.68±1.66 | ns |
| Fractional anisotropy, mean ± SD | 0.22±0.04 | 0.22±0.04 | ns | 0.27±0.04 | 0.27±0.03 | ns |
| Postnatal stress, median [range] | -0.43 [-2.84,2.62] | -0.73 [-2.56, 3.24] | ns | -1.29 [-3.53, 1.42] | -1.48 [-2.93, 1.53] | ns |
| Days of morphine, mean ± SD | 2.68±4.36 | 2.29±2.25 | ns | 2.95±3.99 | 3.20±5.79 | ns |
| Prenatal corticosteroids [yes/no] | 36/5 | 26/4 | ns | 39/3 | 47/2 | ns |
| Postnatal corticosteroids [yes/no] | 12/29 | 9/21 | ns | 15/27 | 17/32 | ns |
| Intraventricular hemorrhaging (yes/no) | 15/26 | 9/21 | ns | 12/30 | 11/38 | ns |
| Necrotizing enterocolitis [yes/no] | 5/36 | 5/25 | ns | 1/41 | 4/45 | ns |
| Retinopathy of prematurity [yes/no] | 15/26 | 13/17 | ns | 17/25 | 13/36 | ns |
| Internalizing symptoms T-score,<br>median [range] | 43 [33-55] | 58 [49-73] | < 0.001 | 41 [29-51] | 58 [47-74] | < 0.001 |

<sup>a</sup> Dutch Perinatal registry reference data (Perined)
Statistical significance was assessed with either a T-test (for continuous data) or a Kruskal-Wallis test (for ordinal data).

**Figure 2.**
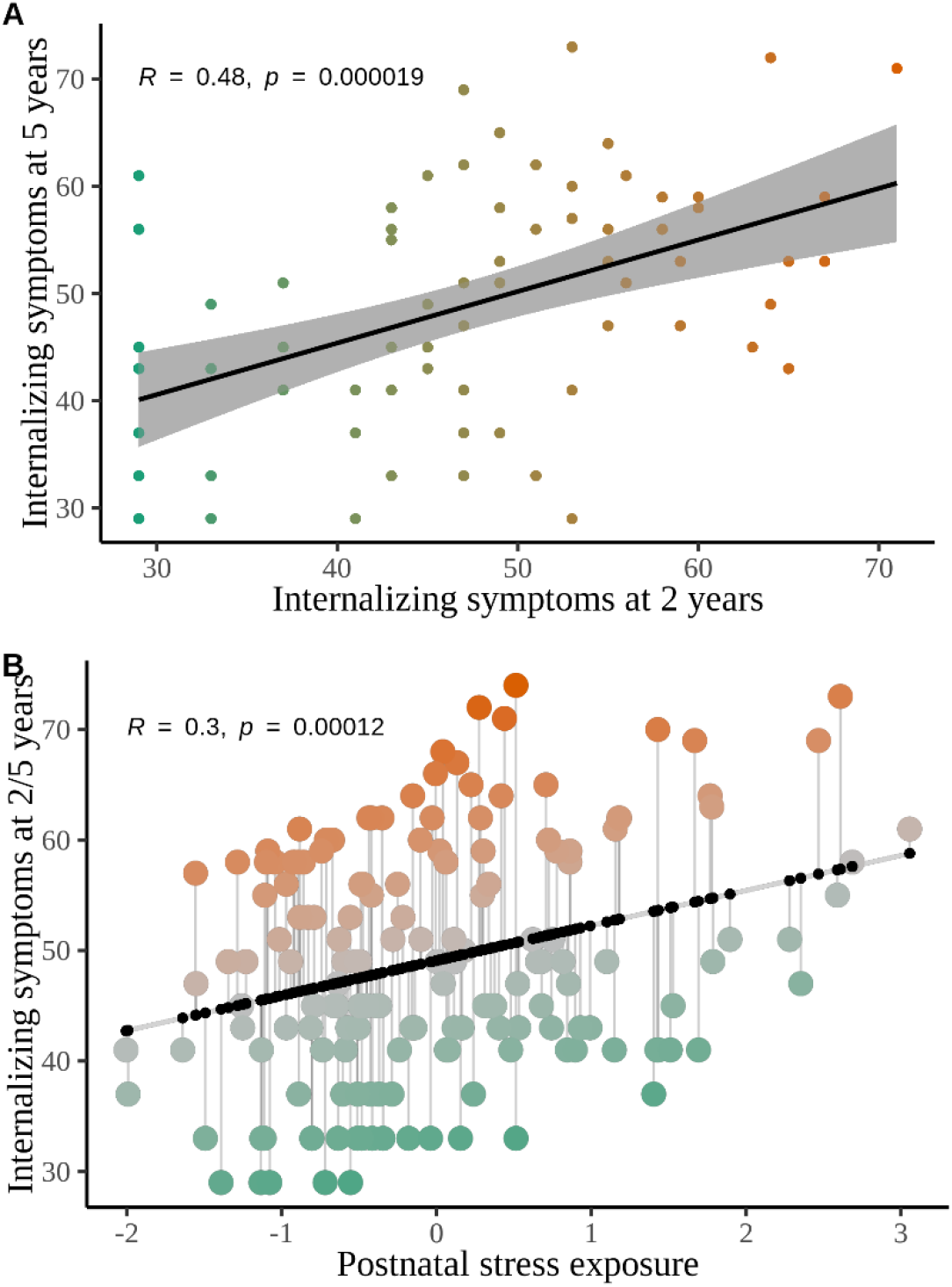
**A**. Significant and positive association between internalizing symptoms assessed at 2 and 5 years of age. **B**. Residualisation approach; orange observations are categorized as stress-overreactve (vulnerable), and green observations are characterized as stress-underreactive (resilient).

Resilience was quantified as a metric of mental health by indexing the internalizing symptoms subscale of the CBCL, taking into account the degree of NICU-related stressor exposure using simple linear regression. We observed a significant positive association between postnatal stress and early childhood internalizing symptoms (*t*(11,151) = 4.08, *p* < 0.001). The fitted regression line (see Figure 2-B) reflected the normative level, with participants positioned above the linear line (i.e., positive residual) expressing an over-reactivity of behavioral symptoms to stressor exposure in the neonatal period and data points below the linear line (i.e., negative residual) representing individuals with under-reactivity to stressor exposure (Amstadter et al., 2014; Van Harmelen et al., 2017).

Preterm-born individuals were classified accordingly: the resilient group showed fewer behavioral symptoms than expected, and a vulnerable group showed more behavioral symptoms problems than expected.

### Statistical analysis

Analyses (connectome development and group-differences, see below) were corrected for confounding factors, including gender, birthweight (z-scores), mean FA, gestational age, age at scan, degree of brain injury (i.e., intraventricular hemorrhage), neonatal surgeries, administration of pre-and postnatal corticosteroids (i.e., accelerates lung maturation), and days of morphine.

#### Stress and connectome development

Longitudinal changes in whole-brain structural connectivity between 30 and 40 weeks of gestation were examined using a time×postnatal stress interaction model using network-based statistic (NBS), a permutation-based method specifically designed to statistically assess network differences (Zalesky et al., 2010). We created a NBS linear-mixed model adjusting for gender, gestational age, age at scan, degree of brain injury (i.e., intraventricular hemorrhage), surgeries, administration of pre-and postnatal corticosteroids (i.e., accelerates lung maturation), and administration of morphine in days, which was applied to all non-zero N_i_×N_j_ connections of the individual networks (lower triangle; consensus-based threshold). The N×N matrix of F-statistics and matching *p*-values associated with the interaction effect was thresholded at a *p*-value of *p* < 0.05. NBS defines the largest connected component, and the size of the largest component is tested against a null-model of permuting subject labels 10000 times. The subsequent null distribution was used to calculate a *p*-value for the largest identified component. We used the main sample and validated the findings in a separate, independent population (see Table 1).

#### Group-differences between resilient and vulnerable individuals

Differences in network organization between resilient and vulnerable individuals were assessed by examining global and local network metrics from the individual structural matrices at term-equivalent age (R packages *igraph*, *braingraph*; R Core Team, 2021). A GLM was specified to test for significant group-difference in network metrics and is compared to permuted data (on graph- or vertex-level), building a null-distribution. Graph-level analyses were permuted 10000 times and vertex-level measures were permuted 5000 times. To correct for multiple comparison the contrast was thresholded on *p* < 0.001.

Local graph parameters, including clustering coefficient, nodal efficiency, eigenvector centrality, and communicability, were calculated to capture the influence of a region on the network. Global measures included clustering coefficient, modularity, strength, and global efficiency. *Clustering coefficient* describes the tendency of regions to cluster together in triangles and is computed by the ratio between the number of connections between region *i* and its neighbor regions and the total number of possible connections with neighbors. A higher clustering coefficient is considered to be a measure of local network segregation (Rubinov & Sporns, 2010). The global measure is computed by taking the mean clustering coefficient of all individual regions in the network. *Nodal efficiency* describes for every region in the network the length of the shortest paths between a given region *i* and all other regions *j*, and measures the average lengths of all shortest paths identified for region *i* (Achard & Bullmore, 2007). Higher nodal efficiency is indicative of a higher capability of information integration, and these regions can also be categorized as a hub. The global measure is computed by taking the mean of nodal efficiency of all individual regions in the network. *Betweenness centrality* describes the influence of a region in the communication between pairs of regions and is measured by the frequency with which a region falls between pairs of other regions on their shortest interconnecting path (Rubinov & Sporns, 2010). This measure reflects the potential influence of a region to control information flow between non-directly connected regions. *Communicability* describes how well a region communicates with every other region in the network and is computed by the weighted sum of all paths and walks between region *i* and *j* (Estrada & Hatano, 2008). High communicability indicates that there are multiple and strong alternative paths connecting the region with other regions. *Modularity* describes the degree to which a network can be organized into modules of densely interconnected regions but sparsely connected between modules and is computed by the difference between the number of edges that lie within a community and a random network of the same degree sequence (Rubinov & Sporns, 2010). High modularity reflects a highly segregated network. *Strength* describes the total sum of the weights of all individual nodal connections in the network. Together, these provide a good understanding of the connectivity and influence of a particular region on the network.

#### Multiclass prediction classification

Random-forest regression with conditional inference trees (RFR-CIF) was used to assess how well node-wise centrality measures could predict the correct classification of the resilient (stress-underreactive) and vulnerable (stress-overreactive) individuals. The predictive multiclass model consisted of a centrality measure (i.e., betweenness centrality) of 47 grey-matter nodes. Analyses were repeated using the other node-wise centrality measures (see Table 4). The predictors were used to build and validate a predictive multiclass model that best fit the combined (main and validation) dataset using 10-fold cross-validation and was tested using a hold-out dataset (65% build and validation [n = 105], 35% testing [n = 57]). The model was fitted on the combined main and validation dataset to increase reliability in estimating probabilities. Slight differences in features due to technical variability in acquisition protocol were removed while preserving biological variability using ComBat prior to model fitting (Fortin et al., 2017, 2018; Johnson et al., 2007).

## Results

The sample consisted of a main (*N*=145, *M*_age_=26.53, *M*_sd_=0.97, 43.5% female) and validation (*N*=123, *M*_age_=26.54, *M*_sd_=1.00, 39.8% female) dataset of preterm born individuals. Both the main (*n*=71) and validation (*n*=91) dataset have follow-up data on parent-reported internalizing symptoms. Key demographics of the two samples are presented in Table 1.

### The effects of postnatal stress on the development of whole-brain structural connectivity

We performed network-based statistics (NBS; see Methods for details) to identify sub-networks of edge-wise effects that showed significant alterations in growth depending on the degree of postnatal stress exposure. NBS analysis revealed one significant cluster of connections, involving 48 connections, with slower growth in connectivity strength from 30 to 40 weeks of gestation for individuals exposed to higher stress (*p* = 0.003, consensus-based threshold, see Figure 1A and 2A). The cluster spanned both hemispheres, involving 20 brain regions such as the amygdala, thalamus, caudate nucleus, and cortical regions such as the insula, fusiform, parahippocampal gyrus, anterior/posterior cingulate cortex, parietal lobe, and frontal lobe. Figure 3 provides a matrix of the vertices and edges involved. The sub-network reduced in size but remained significant across prevalence thresholds (Figure 4E). Also, postnatal stress significantly affected white-matter connectivity at term-equivalent age, with higher stress resulting in lower structural connectivity in a sub-network of 49 connections (Figure 4C, *p* = 0.014).

**Figure 3.**
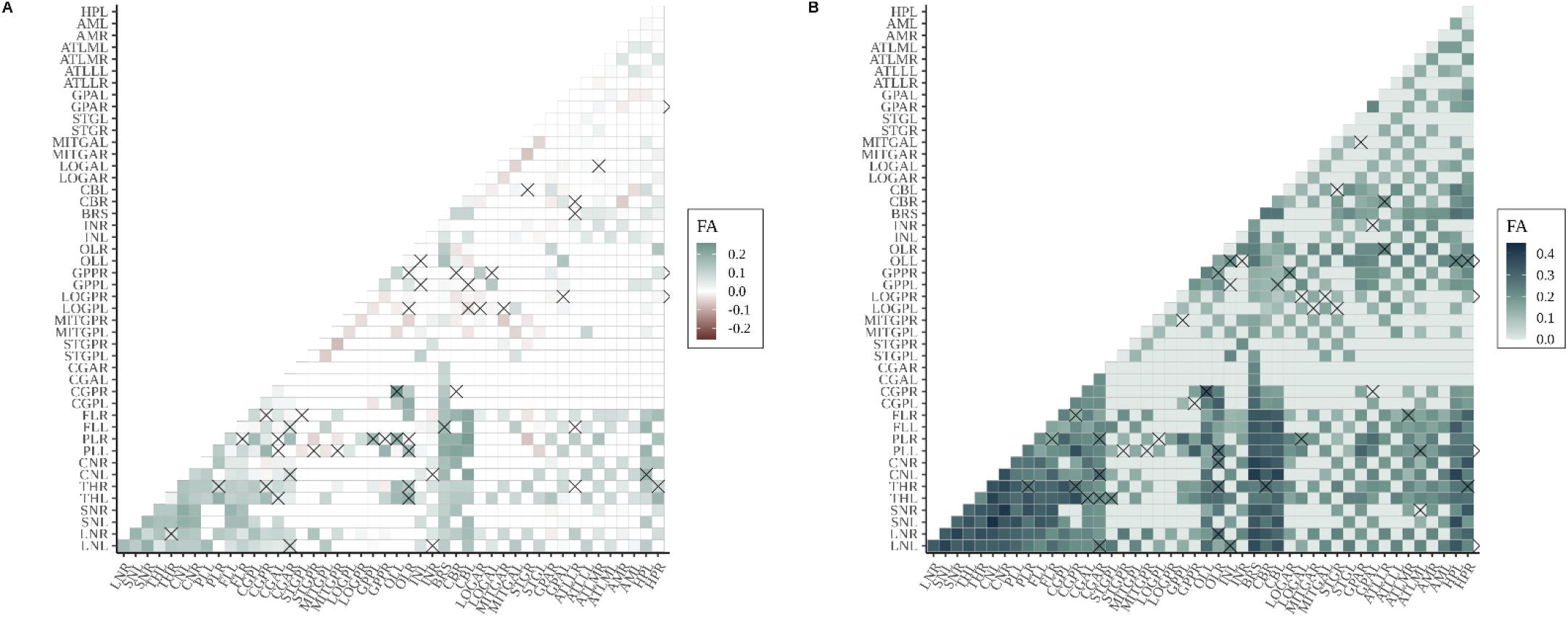
Matrix of largest significant subnetwork (NBS threshold of 50%) identified in the main dataset. Edges with a cross are part of the subnetwork showing a significant time×stress effect (matrix shows the delta in mean connectivity between 30 and 40 weeks of gestation) (**A**) or a significant main effect of stress (**B**) at term-equivalent age. An overview of abbreviations can be found in Figure 1. FA = fractional anisotropy.

**Figure 4.**
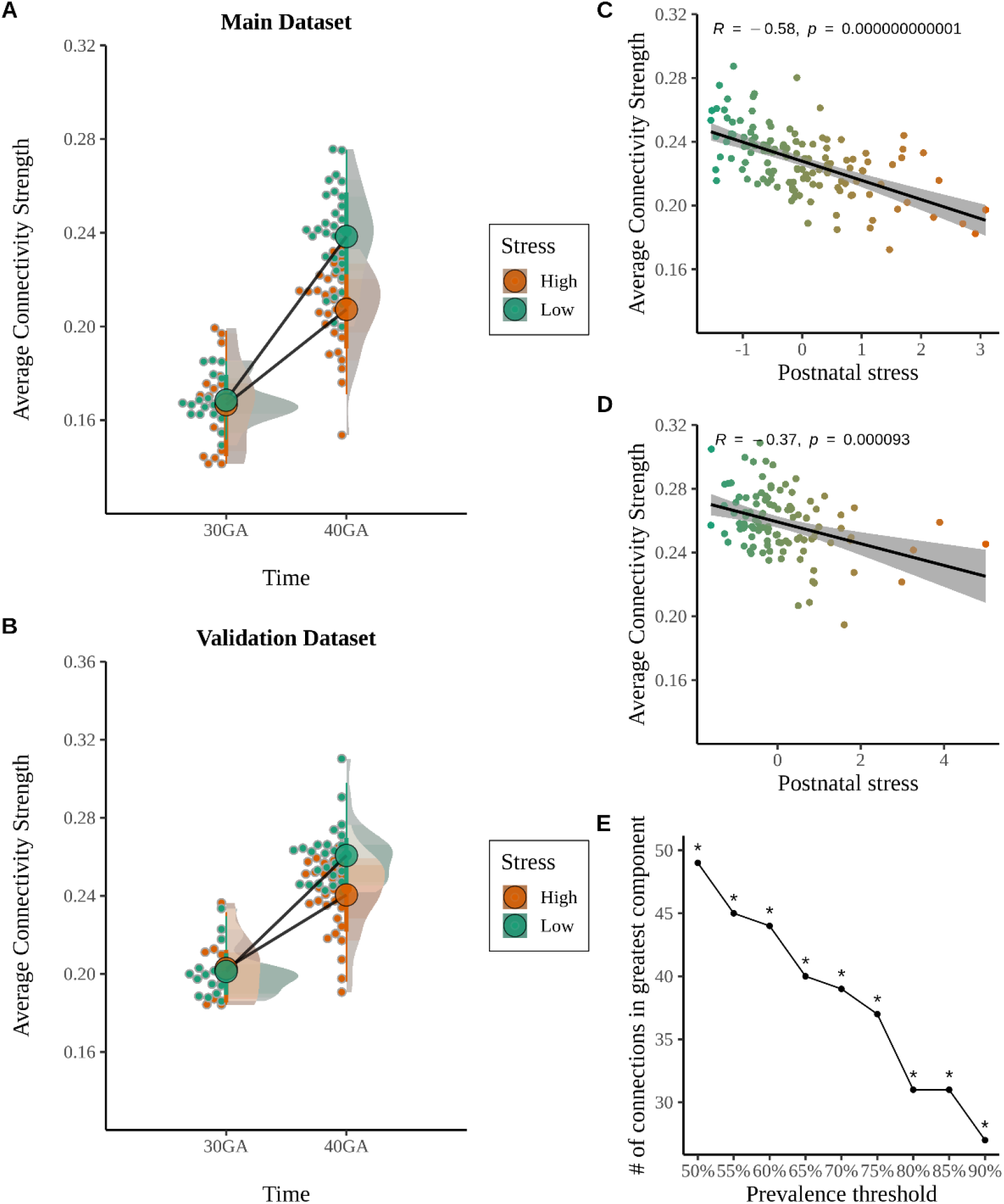
**A**. Schematic representation of the longitudinal time×stress effects, orange representing high stress (highest 25%) and green representing low stress (lowest 25%). **B**. Replication of time×stress effects in an independent sample (32 directions diffusion protocol). **C**. Negative effect of postnatal stress on structural connectivity at term-equivalent age (included 49 connections; 45 directions diffusion protocol). **D**. Replication of stress effects in an independent sample (32 directions diffusion protocol). **E**. Robustness of NBS findings across a range of prevalence thresholds (50%: *p* < 0.05; prevalence threshold of 60%: *p* < 0.05, prevalence threshold of 70%: *p* < 0.05, two-sided permutation testing, 10,000 permutations).

The NBS findings were replicated in an independent sample, providing robust evidence for the effects of postnatal stress on the growth of white-matter connectivity. We masked the connectivity matrix such that only connections were retained if they were part of the sub-network identified in the main sample. Then, we calculated a non-zero mean of connectivity strength and tested the effects of postnatal stress on changes in connectivity strength between 30 and 40 weeks of gestation. We observed a significant stress×time interaction such that higher levels of postnatal stress were associated with slower growth in connectivity strength (Estimate=−0.007(0.003), *F*(1, 37) = 4.79, *p* = 0.035, 95% CI [−0.014, −0.001], see Figure 4B). Also, higher stress was associated with significantly lower levels of white-matter connectivity at term-equivalent age (*t*(13,96) = − 2.44, *p* = 0.016, see Figure 4D).

### Network architecture at term-equivalent age reveal differences between resilient and vulnerable individuals

Based on the normative levels of stress-reactivity (based on the relationship between postnatal NICU-related stress and long-term behavioral symptoms, see “*Resilience to postnatal stress*” Methods), 41 and 42 neonates were classified as stress under-reactive (now being referred to as resilient), and 30 and 49 infants were classified as stress over-reactive (now being referred to as vulnerable). There were no group differences in birth weight, age at birth, age at scan, corticosteroids, days of morphine administration, and mean FA (see Table 2). There was, however, a slight difference in gender in the main dataset (included as a covariate). The reported findings below were thresholded on 75% prevalence, i.e., connections were included if they were reported in at least 75% of the participants. The results reported below are based on structural connectivity at term-equivalent age.

#### Global measures

Analyses revealed no significant group effects in measures of global network architecture.

#### Local measures

We observed significant group effects on local network measures. Group differences were region-specific such that both reduced and increased centrality were observed in vulnerable relative to resilient individuals (see Table 3).

**Table 3.**
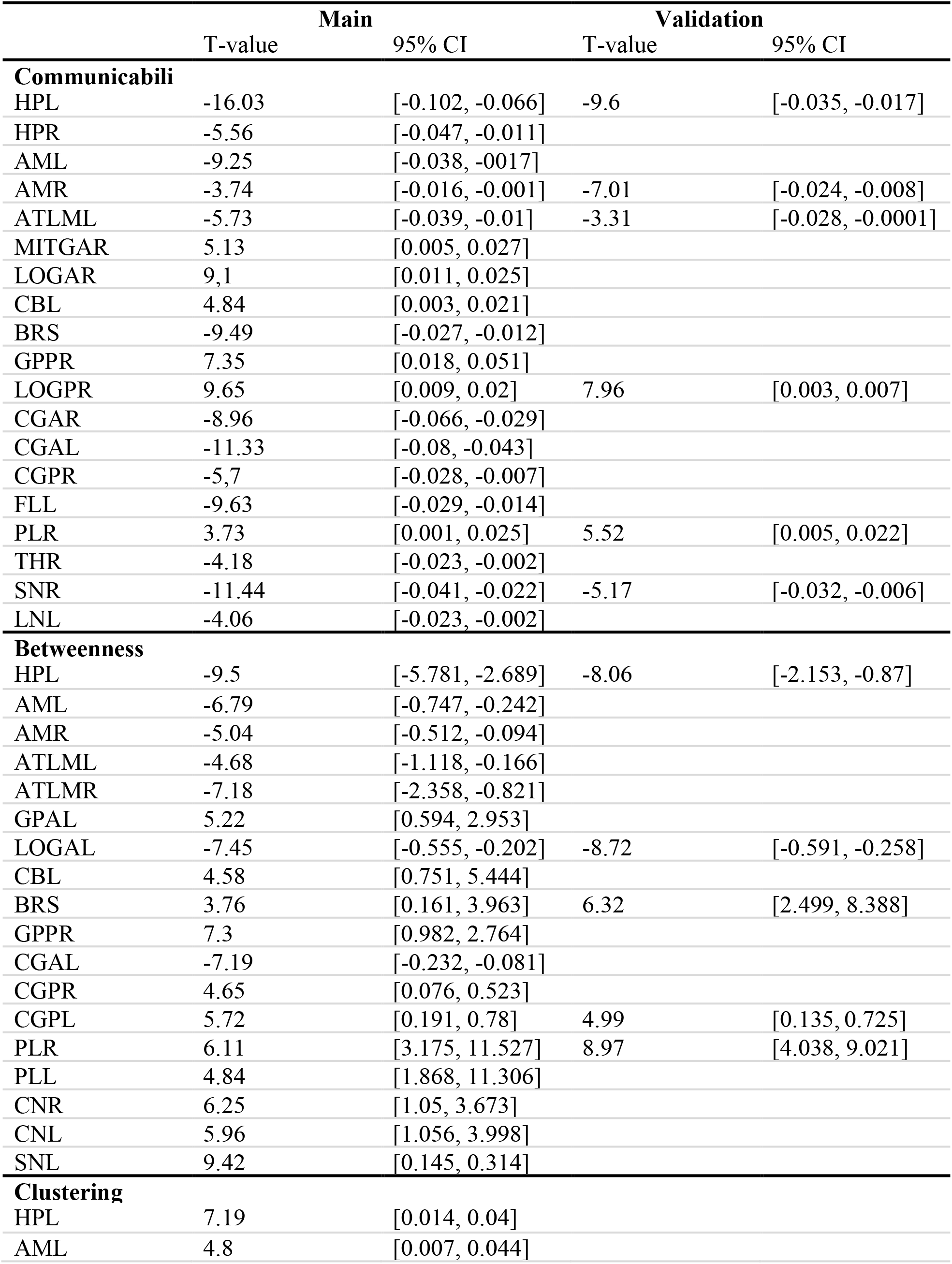
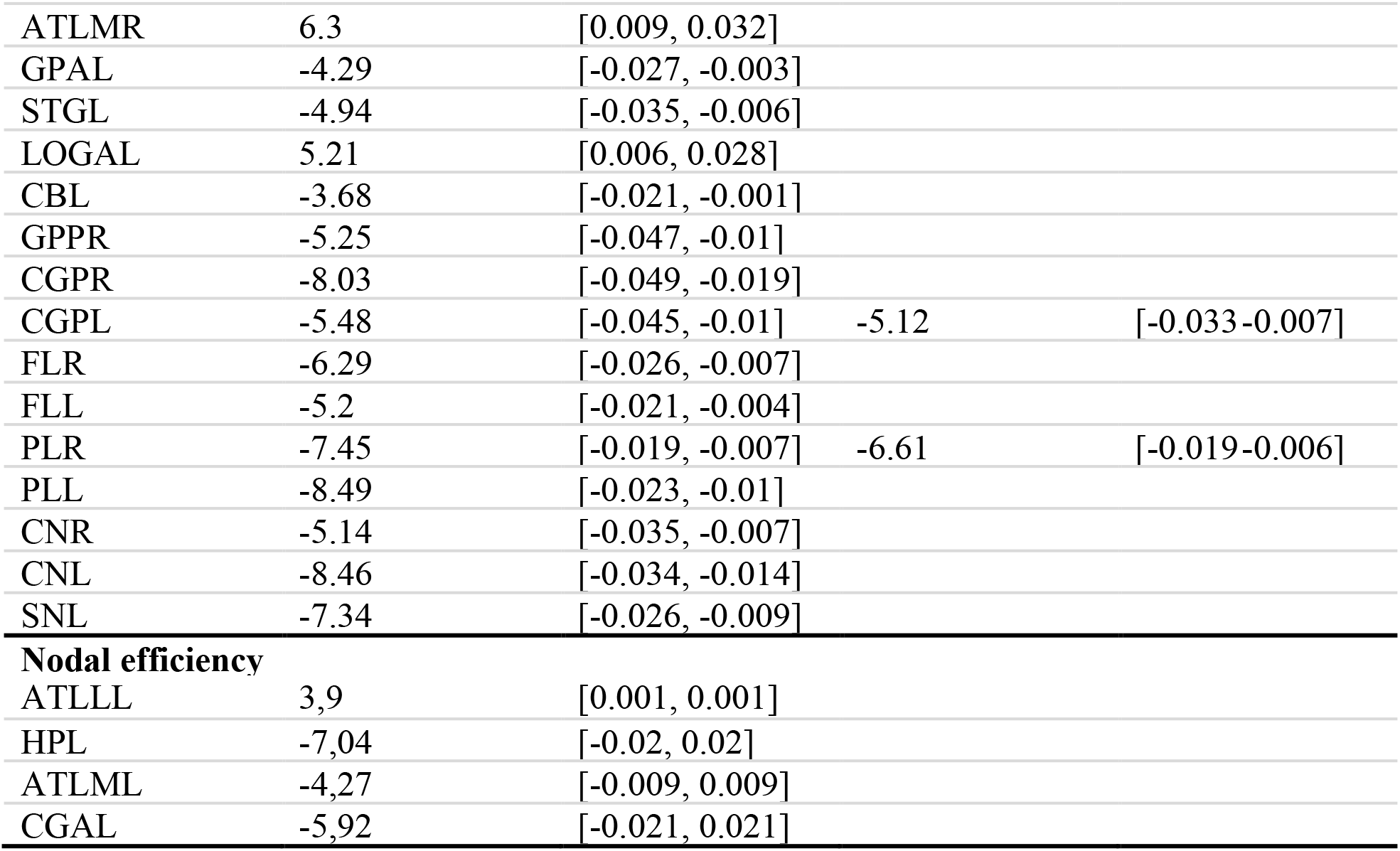
Group-difference on nodal centrality measures for contrast vulnerable > resilient.

|  | **Main** |  | **Validation** |  |
| --- | --- | --- | --- | --- |
|  | T-value | 95% CI | T-value | 95% CI |
| **Communicability** |  |  |  |  |
| HPL | -16.03 | [-0.102, -0.066] | -9.6 | [-0.035, -0.017] |
| HPR | -5.56 | [-0.047, -0.011] |  |  |
| AML | -9.25 | [-0.038, -0.017] |  |  |
| AMR | -3.74 | [-0.016, -0.001] | -7.01 | [-0.024, -0.008] |
| ATLML | -5.73 | [-0.039, -0.01] | -3.31 | [-0.028, -0.0001] |
| MITGAR | 5.13 | [0.005, 0.027] |  |  |
| LOGAR | 9.1 | [0.011, 0.025] |  |  |
| CBL | 4.84 | [0.003, 0.021] |  |  |
| BRS | -9.49 | [-0.027, -0.012] |  |  |
| GPPR | 7.35 | [0.018, 0.051] |  |  |
| LOGPR | 9.65 | [0.009, 0.02] | 7.96 | [0.003, 0.007] |
| CGAR | -8.96 | [-0.066, -0.029] |  |  |
| CGAL | -11.33 | [-0.08, -0.043] |  |  |
| CGPR | -5.7 | [-0.028, -0.007] |  |  |
| FLL | -9.63 | [-0.029, -0.014] |  |  |
| PLR | 3.73 | [0.001, 0.025] | 5.52 | [0.005, 0.022] |
| THR | -4.18 | [-0.023, -0.002] |  |  |
| SNR | -11.44 | [-0.041, -0.022] | -5.17 | [-0.032, -0.006] |
| LNL | -4.06 | [-0.023, -0.002] |  |  |
| **Betweenness** |  |  |  |  |
| HPL | -9.5 | [-5.781, -2.689] | -8.06 | [-2.153, -0.87] |
| AML | -6.79 | [-0.747, -0.242] |  |  |
| AMR | -5.04 | [-0.512, -0.094] |  |  |
| ATLML | -4.68 | [-1.118, -0.166] |  |  |
| ATLMR | -7.18 | [-2.358, -0.821] |  |  |
| GPAL | 5.22 | [0.594, 2.953] |  |  |
| LOGAL | -7.45 | [-0.555, -0.202] | -8.72 | [-0.591, -0.258] |
| CBL | 4.58 | [0.751, 5.444] |  |  |
| BRS | 3.76 | [0.161, 3.963] | 6.32 | [2.499, 8.388] |
| GPPR | 7.3 | [0.982, 2.764] |  |  |
| CGAL | -7.19 | [-0.232, -0.081] |  |  |
| CGPR | 4.65 | [0.076, 0.523] |  |  |
| CGPL | 5.72 | [0.191, 0.78] | 4.99 | [0.135, 0.725] |
| PLR | 6.11 | [3.175, 11.527] | 8.97 | [4.038, 9.021] |
| PLL | 4.84 | [1.868, 11.306] |  |  |
| CNR | 6.25 | [1.05, 3.673] |  |  |
| CNL | 5.96 | [1.056, 3.998] |  |  |
| SNL | 9.42 | [0.145, 0.314] |  |  |
| **Clustering** |  |  |  |  |
| HPL | 7.19 | [0.014, 0.04] |  |  |
| AML | 4.8 | [0.007, 0.044] |  |  |
| ATLMR | 6.3 | [0.009, 0.032] |  |  |
| GPAL | -4.29 | [-0.027, -0.003] |  |  |
| STGL | -4.94 | [-0.035, -0.006] |  |  |
| LOGAL | 5.21 | [0.006, 0.028] |  |  |
| CBL | -3.68 | [-0.021, -0.001] |  |  |
| GPPR | -5.25 | [-0.047, -0.01] |  |  |
| CGPR | -8.03 | [-0.049, -0.019] |  |  |
| CGPL | -5.48 | [-0.045, -0.01] | -5.12 | [-0.033-0.007] |
| FLR | -6.29 | [-0.026, -0.007] |  |  |
| FLL | -5.2 | [-0.021, -0.004] |  |  |
| PLR | -7.45 | [-0.019, -0.007] | -6.61 | [-0.019-0.006] |
| PLL | -8.49 | [-0.023, -0.01] |  |  |
| CNR | -5.14 | [-0.035, -0.007] |  |  |
| CNL | -8.46 | [-0.034, -0.014] |  |  |
| SNL | -7.34 | [-0.026, -0.009] |  |  |
| **Nodal efficiency** |  |  |  |  |
| ATLLL | 3,9 | [0.001, 0.001] |  |  |
| HPL | -7,04 | [-0.02, 0.02] |  |  |
| ATLML | -4,27 | [-0.009, 0.009] |  |  |
| CGAL | -5,92 | [-0.021, 0.021] |  |  |

We first examined the contribution of regions in local network organization as measured by ‘nodal clustering’. Vulnerable infants, relative to resilient, showed a lower clustering of several cortical brain regions overall, including the posterior cingulate cortex (*t*(69) = −5.48, *p* < 0.001), parahippocampal gyrus (*t*(69) = −5.25, *p* < 0.001), frontal lobe (*t*(69) = −6.29, p = *p* < 0.001), and parietal lobe (*t*(69) = −6.61, p = *p* < 0.001). In contrast, higher clustering was observed in the hippocampus (*t*(69) = 7.19, p = *p* < 0.001), amygdala (t(69) = 4.8, p = *p* < 0.001), and medial anterior temporal lobe (*t*(69) = 6.3, p = *p* < 0.001). It is important to note that only differences in the posterior cingulate cortex and parietal lobe were successfully replicated in the validation sample. Statistical details of group differences found in the main and validation dataset can be found in Table 3.

We assessed the contribution of regions in global communication across the brain through ‘betweenness centrality’. On average, vulnerable infants showed a lower centrality of the hippocampus (*t*(69) = −9.5, p = *p* < 0.001) and the anterior fusiform (*t*(69) = −7.45, p = *p* < 0.001), whereas a higher centrality was observed in the brain stem (*t*(69) = 3.76, p = *p* < 0.001), posterior cingulate cortex (*t*(69) = 5.72, p = *p* < 0.001), and parietal lobe (*t*(69) = 6.11, p = *p* < 0.001, see Table 3). These results suggest differential susceptibility in connections central to global brain communication.

We further examined global network integration through ‘communicability’, a metric that considers all possible communication paths between regions in the network. Vulnerable individuals showed, on average, lower communicability of the hippocampus (*t*(69) = −16.03, *p* < 0.001), amygdala (*t*(69) = −3.74, *p* < 0.001), and subthalamic nucleus (*t*(69) = −11.44, *p* < 0.001, see Figure 5 and Table 3). A higher global integration was observed in the posterior parahippocampal gyrus (*t*(69) = 9.65, *p* < 0.001), posterior fusiform (*t*(69) = 9.65, *p* < 0.001), and parietal lobe (*t*(69) = 3.73, *p* < 0.001).

**Figure 5.**
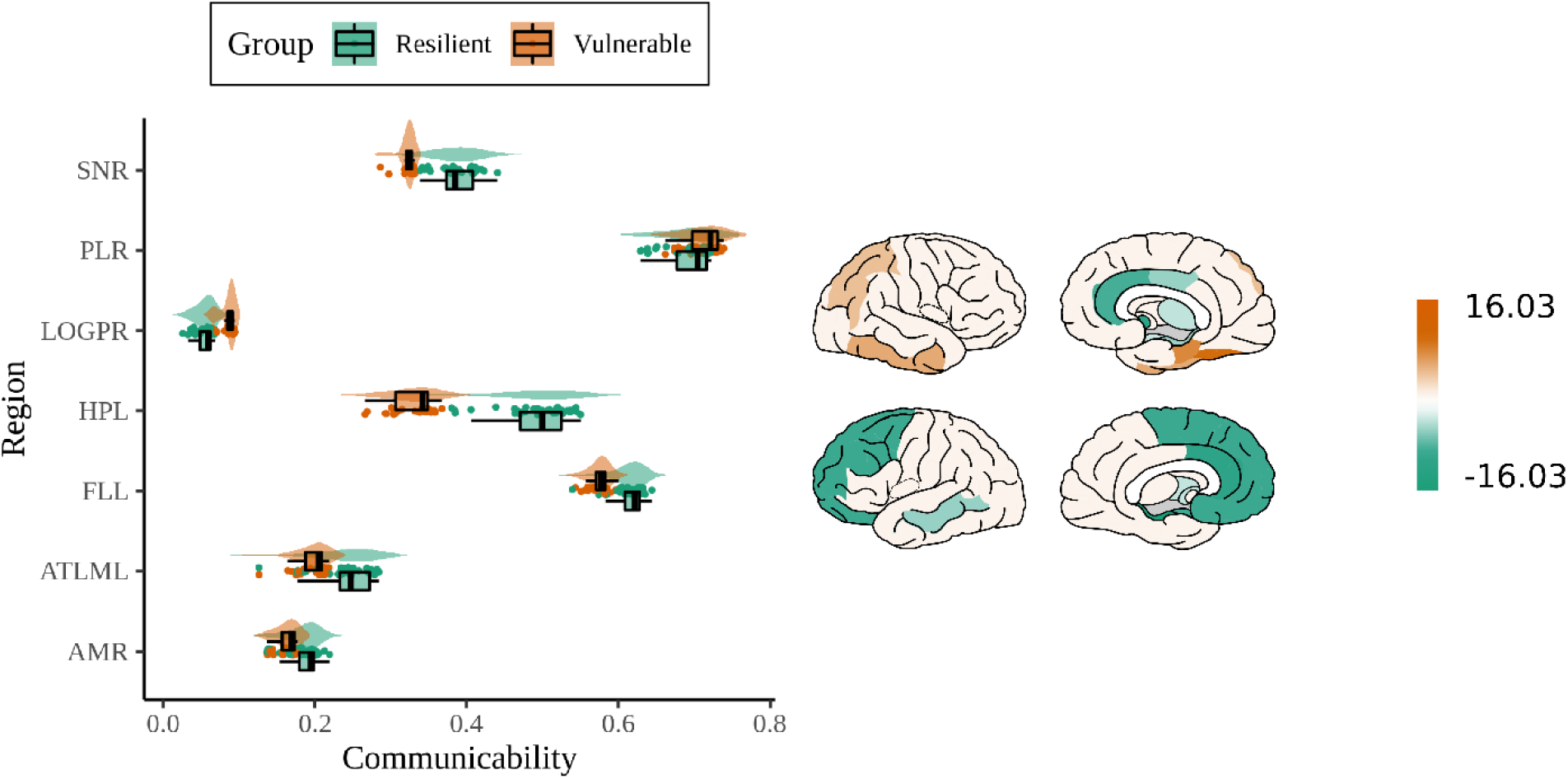
Distribution of group differences in communicability values between vulnerable and resilient infants (left) and regions colored according to T-value (right; vulnerable < resilient [green], Scholtens, L. H, de Lange, S. C., & van den Heuvel, 2021).

Resilient and vulnerable infants did not differ on measures of nodal efficiency.

### Multiclass predictive classification

Random Forest regression with conditional inference trees was used to investigate potential predictive power from local network metrics. Local network measures (i.e., communicability) of the 47 (sub-)cortical grey matter regions were able to correctly classify vulnerable and resilient individuals with an accuracy of 80.4% (*p* < 10^−5^, κ = 0.606, AUC = 0.914). The combined sample (i.e., main and validation) correctly identified the groups with better than 80% balanced accuracy (see Table 4). Importantly, similar results were obtained with the other centrality measures. For model classification and calibration, see Figure 6.

**Table 4.** Multiclass classification using 10-fold cross-validation

|  | Vulnerable<br>versus Resilient | Overall |
| --- | --- | --- |
| **Communicability** |  |  |
| Sensitivity | 0.778 | Accuracy: 0.804 |
| Specificity | 0.828 | 95% CI: [0.676, 0.898] |
| Balanced accuracy | 0.803 | K=0.606 |
| **Betweenness centrality** |  |  |
| Sensitivity | 0.704 | Accuracy: 0.768 |
| Specificity | 0.828 | 95% CI: [0.636, 0.87] |
| Balanced accuracy | 0.766 | K =0.533 |
| **Nodal efficiency** |  |  |
| Sensitivity | 0.630 | Accuracy: 0.746 |
| Specificity | 0.862 | 95% CI: [0.616-0.856] |
| Balanced accuracy | 0.75 | K =0.495 |
| **Clustering coefficient** |  |  |
| Sensitivity | 0.741 | Accuracy: 0.77 |
| Specificity | 0.793 | 95% CI: [0.636-0.87] |
| Balanced accuracy | 0.767 | K =0.534 |

**Figure 6.**
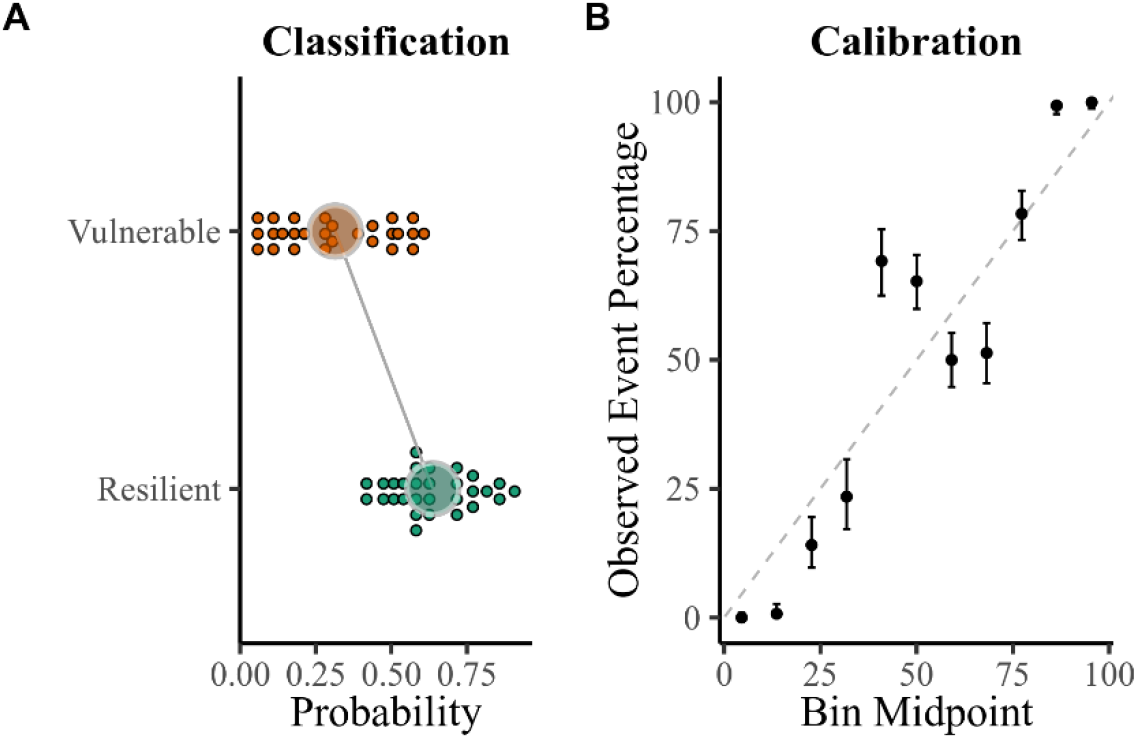
**A**. Shows the difference between the mean of predicted probabilities of the vulnerable an resileitn group. **B**. Shows the true frequency of the positive label against its predicted probability, the x-axis represents the average predicted probability in each bin and the y-axis is the proportion of sample whose class is the vulnerable class (*fraction of positives*).

## Discussion

Preterm-born infants have a life-long increased risk for stress-related psychopathology characterized by anxiety and socio-emotional problems (Arpi & Ferrari, 2013; Upadhyaya et al., 2021). The current study showed that higher stress exposure during NICU admission is associated with slower growth in regions such as the amygdala, hippocampus, insula, and posterior cingulate cortex. Despite these global alterations in development, resilient infants at term-equivalent age can propagate information through regions central for bottom-up emotion regulation. We observed an excellent predictive accuracy of group membership using local network measures at term-equivalent age shortly following exposure. The extra-uterine, postnatal, stressful environment contributes to significant alterations in brain development, but only a proportion of infants show a higher susceptibility for future behavioral problems. A developmental approach is needed to understand longitudinal brain growth following postnatal stress and the neurobiological mechanisms that might confer resilience or vulnerability later in life.

Our findings underscore the impact of postnatal stress on the growth of structural brain connections in corticolimbic pathways across both hemispheres, including critical regions involved in (bottom-up) emotion regulation and processing such as the amygdala, insula, hippocampus, parahippocampal gyrus, and posterior cingulate cortex. These findings align with evidence from other neuroimaging studies showing a delayed development in white matter pathways following preterm birth relative to term-controls (Bouyssi-Kobar et al., 2018; Dodson et al., 2017; Duerden et al., 2018). We now show evidence that in addition to the effects of prematurity, stressful early exposure significantly contributed to a more pronounced impact on delayed development in a sub-network of connections. Interestingly, our findings indicate that resilient individuals can compensate for global alterations in white-matter pathways or reconfigure the brain’s large-scale architecture by selecting resources facilitating information flow throughout the network.

Measures of integration estimate the efficiency of communication among all nodes in a network, enabling the integration and distribution of neural information between spatially distant brain regions (Rubinov & Sporns, 2010). Vulnerable individuals showed lower global integration of the hippocampus, a region that is a crucial regulator of the hypothalamic-pituitary-adrenal axis activation and plays a critical role in the storage and retrieval of emotional memories (Chan et al., 2014; Duval et al., 2015). Prior studies on (early-life) trauma indicated that the hippocampus is particularly vulnerable to chronic pain and stress, with lower volumes and a hypoconnectivity following early-life trauma (Andersen et al., 2008; Shin & Liberzon, 2009). We observed a similar pattern for the amygdala and subthalamic nucleus. The amygdala is part of the (medial) temporal lobe and densely connected with the prefrontal cortex, and has extensive anatomical connections with the paraventricular thalamus and hippocampus. This region plays a critical role in perception, regulation, and plasticity of emotion (Davis & Whalen, 2000; Yang et al., 2017). A less interconnected amygdala in vulnerable infants might seem contradictory, as it does not agree with studies showing evidence of lower amygdala connectivity in resilient trauma-exposed adults (Roeckner et al., 2021). However, a less interconnected amygdala might also be evidence of a decreased inhibitory control of more segregated, cortical, regions including the ventromedial prefrontal cortex (vmPFC) (Andrewes & Jenkins, 2019; Johnstone et al., 2007; Rogers et al., 2017). The lower centrality of the frontal lobe in vulnerable infants substantiates this interpretation. Hence, the increased integration of the hippocampus and amygdala might be a key system in a healthy adaptation with resilience following early disturbances of preterm birth.

Previous studies in children and adults with depression and anhedonia consistently reported a lower capacity for integration (Cullen et al., 2014; Yang et al., 2017). The subthalamic nucleus interconnects with the amygdala and hippocampus and receives convergent cortical and pallidal projections (Accolla et al., 2016) and plays a role in threat appraisal (Serranová et al., 2011). Although the subthalamic nucleus received attention concerning Parkinson’s disease, the increased social and affective alterations following deep-brain stimulation have been implicated in the emergence of enhanced affective processing and decreased depressive symptoms (Schneider et al., 2003; Smeding et al., 2006). Higher integration of these regions might be beneficial in retaining mental wellbeing following preterm birth.

Preterm-born infants (Bouyssi-Kobar et al., 2019; Sa de Almeida et al., 2021) and children (de Kieviet et al., 2021; Young et al., 2018) show alterations in information flow. A balance between integration and segregation is essential for efficient communication through local processing and global communication. A lower integration between regions important for fear memory and emotion-processing in vulnerable infants renders their networks more susceptible. It puts them at increased risk for behavioral problems by making it harder to effectively compensate for second-hit abnormalities that might occur within a network or region.

The dysconnectivity of neuroanatomical networks has been implicated in the emergence of several neurological and psychiatric disorders, including anxiety and depression (Akiki et al., 2018; Sang et al., 2018; Yu et al., 2013). The relative preservation of global integration of regions implicated in emotion processing and regulation may support impaired white matter pathways more effectively after preterm birth. These results might indicate that resilient individuals can increase and diminish information flow of specific regions, enabling them to compensate for global alterations following preterm birth. Interestingly, however, it remains elusive whether a higher centrality of the hippocampus could be interpreted as a compensatory adaptation following preterm birth, enabling preterm-born individuals to “bounce back” or if it could be considered a preexisting protective factor. Resilience studies following childhood trauma indicated that an increased hippocampal connectivity aids emotion regulation and enables one to successfully cope with trauma (Richter et al., 2019; van Rooij et al., 2021). Similarly, resilient individuals exhibit lower insula activity, facilitating an appropriate adjustment of emotional resources (Haase et al., 2016; Waugh et al., 2008). Although the current study contributes to the literature of resilience following preterm birth, future studies investigating the (neuroprotective) mechanisms by which global integration is increased in resilient infants are warranted.

While a direct inverse relationship between resilience and vulnerability does not exist, vulnerability studies seem to present the best available approximation for the concept of resilience in preterm-born individuals. In line with studies on trauma exposure, preterm-born individuals with more problem behavior seem to show reduced hippocampal connectivity and lower volumes (Aanes et al., 2015; Rogers et al., 2018) and a lower interconnected amygdala (Rogers et al., 2017). Further, preliminary interventional studies focusing on neuroprotection, reducing the impact of postnatal stress following preterm birth (e.g., music and massage therapy) showed significantly improved white matter maturation of the uncinate fasciculus (Sa de Almeida et al., 2020). Hence, these results implicate that increasing information flow of the amygdala and hippocampus may lead to symptom attenuation and is consistent with our observation that vulnerable and resilient individuals differ in a small number of regions or pathways that may facilitate compensation.

The differences in neural representations between resilience and vulnerable infants enable the accurate classification of group membership. The current study shows that at term-equivalent age, the connectome shows distinguishable features in topological architecture. Demonstrating these patterns highlights that resilience and vulnerability occur in the context of unique neurobiological differentiability and may be considered a valuable biomarker for predicting behavioral symptoms in early childhood.

Several methodological issues should be taken into consideration when interpreting our findings. Although our unique dataset enables us to investigate individual differences in longitudinal white-matter development, we could only model linear change. Studies involving three or more time points (Remer et al., 2017) can fit several slopes, including quadratic, logarithmic, and cubic, facilitating a more nuanced understanding of how postnatal stress affects brain development. For instance, a quadratic growth pattern would mean that the effects of postnatal stress emerge during a specific developmental period and then declines or disappear during a particular period and then reappears later. Despite this methodological limitation, our results nevertheless provide convincing evidence that between 30 and 40 weeks of gestation, postnatal stress significantly reduces linear growth in a sub-network of connections. Another limitation is that the resilient infants might have been healthier than the vulnerable infants. Although the residualisation approach controls for the degree of postnatal stress exposure, resilient infants could still have experienced fewer complications and clinical procedures compared to vulnerable infants. Notably, infants did not differ on a large set of clinical parameters (see Table 2). In other words, resilient preterm-born infants did not endure fewer clinical procedures, and it is unlikely that they were healthier than vulnerable infants.

Our longitudinal findings suggest that postnatal stress leads to sparser brain connectivity after preterm birth. Importantly, alterations in specific brain areas impacting bottom-up emotion regulation render preterm infants resilient to internalizing symptoms later in life. These findings emphasize the detrimental impact of postnatal stress and the relative plasticity of the preterm brain. The current results suggest that resilience appertains to a potential compensatory or innate ability to propagate global information flow, informing future intervention studies on fostering specific nodal changes.

## Acknowledgements

Femke Lammertink was supported by a grant from the Wilhelmina Children’s Hospital (D-17-010007). Martijn P. van den Heuvel was supported by a VIDI (452-16-015) grant from the Netherlands Organization for Scientific Research (NWO), and a European Research Council grant (ERC-2015-CoG 101001062). Erno J. Hermans was supported by a European Research Council grant (ERC-2015-CoG 682591). The content is the sole responsibility of the authors and does not necessarily represent the official views of the funding agencies. The authors declare that the research was conducted in the absence of any commercial or financial relationships that could be construed as a potential conflict of interest.

## References

Aanes, S., Bjuland, K. J., Skranes, J., & Løhaugen, G. C. C. (2015). Memory function and hippocampal volumes in preterm born very-low-birth-weight (VLBW) young adults. NeuroImage, 105, 76–83. https://doi.org/10.1016/j.neuroimage.2014.10.023

Achenbach, T., & Rescorla, L. (2001). Manual for the ASEBA School-Age Forms & Profiles.

Akazawa, K., Chang, L., Yamakawa, R., Hayama, S., Buchthal, S., Alicata, D., Andres, T., Castillo, D., Oishi, K., Skranes, J., Ernst, T., & Oishi, K. (2016). Probabilistic maps of the white matter tracts with known associated functions on the neonatal brain atlas: Application to evaluate longitudinal developmental trajectories in term-born and preterm-born infants. NeuroImage, 128, 167–179. https://doi.org/10.1016/j.neuroimage.2015.12.026

Akiki, T. J., Averill, C. L., Wrocklage, K. M., Scott, J. C., Averill, L. A., Schweinsburg, B., Alexander-Bloch, A., Martini, B., Southwick, S. M., Krystal, J. H., & Abdallah, C. G. (2018). Default mode network abnormalities in posttraumatic stress disorder: A novel network-restricted topology approach. NeuroImage, 176, 489–498. https://doi.org/10.1016/j.neuroimage.2018.05.005

Amstadter, A. B., Myers, J. M., & Kendler, K. S. (2014). Psychiatric resilience: Longitudinal twin study. British Journal of Psychiatry, 205(4), 275–280. https://doi.org/10.1192/bjp.bp.113.130906

Andersson, J. L. R., Jenkinson, M., & Smith, S. (2007). Non-linear registration aka spatial normalisation (Technical). Oxford: FMRIB Centre.

Andersson, J. L. R., & Stamatios, S. N. (2016). An integrated approach to correction for off-resonance effects and subject movement in diffusion MR imaging. NeuroImage, 125, 1063–1078. https://doi.org/10.1016/j.neuroimage.2015.10.019

Ball, G., Aljabar, P., Zebari, S., Tusor, N., Arichi, T., Merchant, N., Robinson, E. C., Ogundipe, E., Rueckert, D., Edwards, A. D., & Counsell, S. J. (2014). Rich-club organization of the newborn human brain. Proceedings of the National Academy of Sciences of the United States of America, 111(20), 7456–7461. https://doi.org/10.1073/pnas.1324118111

Ball, G., Boardman, J. P., Aljabar, P., Pandit, A., Arichi, T., Merchant, N., Rueckert, D., Edwards, A. D., & Counsell, S. J. (2013). The influence of preterm birth on the developing thalamocortical connectome. Cortex, 49(6), 1711–1721. https://doi.org/10.1016/j.cortex.2012.07.006

Ball, G., Boardman, J. P., Rueckert, D., Aljabar, P., Arichi, T., Merchant, N., Gousias, I. S., Edwards, A. D., & Counsell, S. J. (2012). The effect of preterm birth on thalamic and cortical development. Cerebral Cortex, 22(5), 1016–1024. https://doi.org/10.1093/cercor/bhr176

Ball, G., Srinivasan, L., Aljabar, P., Counsell, S. J., Durighel, G., Hajnal, J. V., Rutherford, M. A., & Edwards, A. D. (2013). Development of cortical microstructure in the preterm human brain. Proceedings of the National Academy of Sciences of the United States of America, 110(23), 9541–9546. https://doi.org/10.1073/pnas.1301652110

Batalle, D., Hughes, E. J., Zhang, H., Tournier, J. D., Tusor, N., Aljabar, P., Wali, L., Alexander, D. C., Hajnal, J. V., Nosarti, C., Edwards, A. D., & Counsell, S. J. (2017). Early development of structural networks and the impact of prematurity on brain connectivity. NeuroImage, 149, 379–392. https://doi.org/10.1016/j.neuroimage.2017.01.065

Bouyssi-Kobar, M., Brossard-Racine, M., Jacobs, M., Murnick, J., Chang, T., & Limperopoulos, C. (2018). Regional microstructural organization of the cerebral cortex is affected by preterm birth. NeuroImage: Clinical, 18, 871–880. https://doi.org/10.1016/j.nicl.2018.03.020

Bouyssi-Kobar, M., De Asis-Cruz, J., Murnick, J., Chang, T., & Limperopoulos, C. (2019). Altered Functional Brain Network Integration, Segregation, and Modularity in Infants Born Very Preterm at Term-Equivalent Age. Journal of Pediatrics, 213, 13–21.e1. https://doi.org/10.1016/j.jpeds.2019.06.030

Broyd, S. J., Demanuele, C., Debener, S., Helps, S. K., James, C. J., & Sonuga-Barke, E. J. S. (2009). Default-mode brain dysfunction in mental disorders: A systematic review. Neuroscience and Biobehavioral Reviews, 33(3), 279–296. https://doi.org/10.1016/j.neubiorev.2008.09.002

Brummelte, S., Grunau, R. E., Chau, V., Poskitt, K. J., Brant, R., Vinall, J., Gover, A., Synnes, A. R., & Miller, S. P. (2012). Procedural pain and brain development in premature newborns. Annals of Neurology, 71(3), 385–396. https://doi.org/10.1002/ana.22267

Buff, C., Brinkmann, L., Neumeister, P., Feldker, K., Heitmann, C., Gathmann, B., Andor, T., & Straube, T. (2016). Specifically altered brain responses to threat in generalized anxiety disorder relative to social anxiety disorder and panic disorder. NeuroImage: Clinical, 12, 698–706. https://doi.org/10.1016/j.nicl.2016.09.023

Chan, E., Baumann, O., Bellgrove, M. A., & Mattingley, J. B. (2014). Negative emotional experiences during navigation enhance parahippocampal activity during recall of place information. Journal of Cognitive Neuroscience, 26(1), 154–164. https://doi.org/10.1162/jocn_a_00468

Chen, H. J., Qi, R., Ke, J., Qiu, J., Xu, Q., Zhang, Z., Zhong, Y., Lu, G. M., & Chen, F. (2020). Altered dynamic parahippocampus functional connectivity in patients with post-traumatic stress disorder. World Journal of Biological Psychiatry, 1–10. https://doi.org/10.1080/15622975.2020.1785006

De Asis-Cruz, J., Krishnamurthy, D., Zhao, L., Kapse, K., Vezina, G., Andescavage, N., Quistorff, J., Lopez, C., & Limperopoulos, C. (2020). Association of Prenatal Maternal Anxiety With Fetal Regional Brain Connectivity. JAMA Network Open, 3(12), e2022349. https://doi.org/10.1001/jamanetworkopen.2020.22349

de Kieviet, J. F., Lustenhouwer, R., Königs, M., van Elburg, R. M., Pouwels, P. J. W., & Oosterlaan, J. (2021). Altered structural connectome and motor problems of very preterm born children at school-age. Early Human Development, 152, 105274. https://doi.org/10.1016/j.earlhumdev.2020.105274

Degnan, A. J., Wisnowski, J. L., Choi, S., Ceschin, R., Bhushan, C., Leahy, R. M., Corby, P., Schmithorst, V. J., & Panigrahy, A. (2015). Altered Structural and Functional Connectivity in Late Preterm Preadolescence: An Anatomic Seed-Based Study of Resting State Networks Related to the Posteromedial and Lateral Parietal Cortex. PLOS ONE, 10(6), e0130686. https://doi.org/10.1371/journal.pone.0130686

Disselhoff, V., Jakab, A., Schnider, B., Latal, B., Wehrle, F. M., & Hagmann, C. F. (2020). Inhibition is associated with whole-brain structural brain connectivity on network level in school-aged children born very preterm and at term; Inhibition abilities and structural brain connectivity in prematurity. NeuroImage, 218. https://doi.org/10.1016/j.neuroimage.2020.116937

Dodson, C. K., Travis, K. E., Ben-Shachar, M., & Feldman, H. M. (2017). White matter microstructure of 6-year old children born preterm and full term. NeuroImage: Clinical, 16, 268–275. https://doi.org/10.1016/j.nicl.2017.08.005

Eikenes, L., Løhaugen, G. C., Brubakk, A. M., Skranes, J., & Håberg, A. K. (2011). Young adults born preterm with very low birth weight demonstrate widespread white matter alterations on brain DTI. NeuroImage, 54(3), 1774–1785. https://doi.org/10.1016/j.neuroimage.2010.10.037

Fischi-Gomez, E., Muñoz-Moreno, E., Vasung, L., Griffa, A., Borradori-Tolsa, C., Monnier, M., Lazeyras, F., Thiran, J. P., & Hüppi, P. S. (2016). Brain network characterization of high-risk preterm-born school-age children. NeuroImage: Clinical, 11, 195–209. https://doi.org/10.1016/j.nicl.2016.02.001

Gogolla, N. (2017). The insular cortex. In Current Biology (Vol. 27, Issue 12, pp. R580–R586). Cell Press. https://doi.org/10.1016/j.cub.2017.05.010

Greve, D. N., & Fischl, B. (2009). Accurate and robust brain image alignment using boundary-based registration. NeuroImage, 48(1), 63–72. https://doi.org/10.1016/j.neuroimage.2009.06.060

Groppo, M., Ricci, D., Bassi, L., Merchant, N., Doria, V., Arichi, T., Allsop, J. M., Ramenghi, L., Fox, M. J., Cowan, F. M., Counsell, S. J., & Edwards, A. D. (2014). Development of the optic radiations and visual function after premature birth. Cortex, 56, 30–37. https://doi.org/10.1016/j.cortex.2012.02.008

Haase, L., Stewart, J. L., Youssef, B., May, A. C., Isakovic, S., Simmons, A. N., Johnson, D. C., Potterat, E. G., & Paulus, M. P. (2016). When the brain does not adequately feel the body: Links between low resilience and interoception. Biological Psychology, 113, 37–45. https://doi.org/10.1016/j.biopsycho.2015.11.004

Johns, C. B., Lacadie, C., Vohr, B., Ment, L. R., & Scheinost, D. (2019). Amygdala functional connectivity is associated with social impairments in preterm born young adults. NeuroImage: Clinical, 21, 101626. https://doi.org/10.1016/j.nicl.2018.101626

Karolis, V. R., Froudist-Walsh, S., Brittain, P. J., Kroll, J., Ball, G., Edwards, A. D., Dell’Acqua, F., Williams, S. C., Murray, R. M., & Nosarti, C. (2016). Reinforcement of the Brain’s Rich-Club Architecture Following Early Neurodevelopmental Disruption Caused by Very Preterm Birth. Cerebral Cortex, 26(3), 1322–1335. https://doi.org/10.1093/cercor/bhv305

Kilpatrick, L., & Cahill, L. (2003). Amygdala modulation of parahippocampal and frontal regions during emotionally influenced memory storage. NeuroImage, 20(4), 2091–2099. https://doi.org/10.1016/j.neuroimage.2003.08.006

Lautarescu, A., Pecheva, D., Nosarti, C., Nihouarn, J., Zhang, H., Victor, S., Craig, M., Edwards, A. D., & Counsell, S. J. (2020). Maternal Prenatal Stress Is Associated With Altered Uncinate Fasciculus Microstructure in Premature Neonates. Biological Psychiatry, 87(6), 559–569. https://doi.org/10.1016/j.biopsych.2019.08.010

Leech, R., & Sharp, D. J. (2014). The role of the posterior cingulate cortex in cognition and disease. In Brain (Vol. 137, Issue 1, pp. 12–32). Oxford University Press. https://doi.org/10.1093/brain/awt162

Loe, I. M., Lee, E. S., & Feldman, H. M. (2013). Attention and internalizing behaviors in relation to white matter in children born preterm. Journal of Developmental and Behavioral Pediatrics, 34(3), 156–164. https://doi.org/10.1097/DBP.0b013e3182842122

Marusak, H. A., Etkin, A., & Thomason, M. E. (2015). Disrupted insula-based neural circuit organization and conflict interference in trauma-exposed youth. NeuroImage: Clinical, 8, 516–525. https://doi.org/10.1016/j.nicl.2015.04.007

Menegaux, A., Meng, C., Neitzel, J., Bäuml, J. G., Müller, H. J., Bartmann, P., Wolke, D., Wohlschläger, A. M., Finke, K., & Sorg, C. (2017). Impaired visual short-term memory capacity is distinctively associated with structural connectivity of the posterior thalamic radiation and the splenium of the corpus callosum in preterm-born adults. NeuroImage, 150, 68–76. https://doi.org/10.1016/j.neuroimage.2017.02.017

Menon, V. (2011). Large-scale brain networks and psychopathology: A unifying triple network model. Trends in Cognitive Sciences, 15(10), 483–506. https://doi.org/10.1016/j.tics.2011.08.003

Mullen, K. M., Vohr, B. R., Katz, K. H., Schneider, K. C., Lacadie, C., Hampson, M., Makuch, R. W., Reiss, A. L., Constable, R. T., & Ment, L. R. (2011). Preterm birth results in alterations in neural connectivity at age 16 years. NeuroImage, 54(4), 2563–2570. https://doi.org/10.1016/j.neuroimage.2010.11.019

Papini, C., White, T. P., Montagna, A., Brittain, P. J., Froudist-Walsh, S., Kroll, J., Karolis, V., Simonelli, A., Williams, S. C., Murray, R. M., & Nosarti, C. (2016). Altered resting-state functional connectivity in emotion-processing brain regions in adults who were born very preterm. Psychological Medicine, 46(14), 3025–3039. https://doi.org/10.1017/S0033291716001604

R Core Team (2021). R: A language and environment for statistical computing. R Foundation for Statistical Computing, Vienna, Austria. https://www.R-project.org/.

Remer, J., Croteau-Chonka, E., Dean, D. C., D’Arpino, S., Dirks, H., Whiley, D., & Deoni, S. C. L. (2017). Quantifying cortical development in typically developing toddlers and young children, 1–6 years of age. NeuroImage, 153, 246–261. https://doi.org/10.1016/j.neuroimage.2017.04.010

Richter, A., Krämer, B., Diekhof, E. K., & Gruber, O. (2019). Resilience to adversity is associated with increased activity and connectivity in the VTA and hippocampus. NeuroImage: Clinical, 23, 101920. https://doi.org/10.1016/j.nicl.2019.101920

Rimol, L. M., Botellero, V. L., Bjuland, K. J., Løhaugen, G. C. C., Lydersen, S., Evensen, K. A. I., Brubakk, A. M., Eikenes, L., Indredavik, M. S., Martinussen, M., Yendiki, A., Håberg, A. K., & Skranes, J. (2019). Reduced white matter fractional anisotropy mediates cortical thickening in adults born preterm with very low birthweight. NeuroImage, 188, 217–227. https://doi.org/10.1016/j.neuroimage.2018.11.050

Rogers, C. E., Lean, R. E., Wheelock, M. D., & Smyser, C. D. (2018). Aberrant structural and functional connectivity and neurodevelopmental impairment in preterm children. In Journal of Neurodevelopmental Disorders (Vol. 10, Issue 1, pp. 1–13). BioMed Central Ltd. https://doi.org/10.1186/s11689-018-9253-x

Rogers, C. E., Sylvester, C. M., Mintz, C., Kenley, J. K., Shimony, J. S., Barch, D. M., & Smyser, C. D. (2017). Neonatal Amygdala Functional Connectivity at Rest in Healthy and Preterm Infants and Early Internalizing Symptoms. Journal of the American Academy of Child and Adolescent Psychiatry, 56(2), 157–166. https://doi.org/10.1016/j.jaac.2016.11.005

Sa de Almeida, J., Lordier, L., Zollinger, B., Kunz, N., Bastiani, M., Gui, L., Adam-Darque, A., Borradori-Tolsa, C., Lazeyras, F., & Hüppi, P. S. (2020). Music enhances structural maturation of emotional processing neural pathways in very preterm infants. NeuroImage, 207, 116391. https://doi.org/10.1016/j.neuroimage.2019.116391

Sa de Almeida, J., Meskaldji, D. E., Loukas, S., Lordier, L., Gui, L., Lazeyras, F., & Hüppi, P. S. (2021). Preterm birth leads to impaired rich-club organization and fronto-paralimbic/limbic structural connectivity in newborns. NeuroImage, 225, 117440. https://doi.org/10.1016/j.neuroimage.2020.117440

Sang, L., Chen, L., Wang, L., Zhang, J., Zhang, Y., Li, P., Li, C., & Qiu, M. (2018). Progressively disrupted brain functional connectivity network in subcortical ischemic vascular cognitive impairment patients. Frontiers in Neurology, 9(FEB), 94. https://doi.org/10.3389/fneur.2018.00094

Scheinost, D., Kwon, S. H., Shen, X., Lacadie, C., Schneider, K. C., Dai, F., Ment, L. R., & Constable, R. T. (2016). Preterm birth alters neonatal, functional rich club organization. Brain Structure and Function, 221(6), 3211–3222. https://doi.org/10.1007/s00429-015-1096-6

Scholtens, L. H., de Lange, S. C., and van den Heuvel, M. P. (2021). “Simple Brain Plot”. Zenodo. https://doi.org/10.5281/zenodo.5346593

Sølsnes, A. E., Sripada, K., Yendiki, A., Bjuland, K. J., Østgård, H. F., Aanes, S., Grunewaldt, K. H., Løhaugen, G. C., Eikenes, L., Håberg, A. K., Rimol, L. M., & Skranes, J. (2016). Limited microstructural and connectivity deficits despite subcortical volume reductions in school-aged children born preterm with very low birth weight. NeuroImage, 130, 24–34. https://doi.org/10.1016/j.neuroimage.2015.12.029

Song, L., Mishra, V., Ouyang, M., Peng, Q., Slinger, M., Liu, S., & Huang, H. (2017). Human fetal brain Connectome: Structural network development from middle fetal stage to birth. Frontiers in Neuroscience, 11(OCT). https://doi.org/10.3389/fnins.2017.00561

Teicher, M. H., Anderson, C. M., Ohashi, K., & Polcari, A. (2014). Childhood maltreatment: Altered network centrality of cingulate, precuneus, temporal pole and insula. Biological Psychiatry, 76(4), 297–305. https://doi.org/10.1016/j.biopsych.2013.09.016

Turk, E., van den Heuvel, M. I., Benders, M. J., de Heus, R., Franx, A., Manning, J. H., Hect, J. L., Hernandez-Andrade, E., Hassan, S. S., Romero, R., Kahn, R. S., Thomason, M. E., & van den Heuvel, M. P. (2019). Functional Connectome of the Fetal Brain. The Journal of Neuroscience : The Official Journal of the Society for Neuroscience, 39(49), 9716–9724. https://doi.org/10.1523/JNEUROSCI.2891-18.2019

Van Harmelen, A. L., Kievit, R. A., Ioannidis, K., Neufeld, S., Jones, P. B., Bullmore, E., Dolan, R., Fonagy, P., & Goodyer, I. (2017). Adolescent friendships predict later resilient functioning across psychosocial domains in a healthy community cohort. Psychological Medicine, 47(13), 2312–2322. https://doi.org/10.1017/S0033291717000836

van Rooij, S. J. H., Ravi, M., Ely, T. D., Michopoulos, V., Winters, S. J., Shin, J., Marin, M. F., Milad, M. R., Rothbaum, B. O., Ressler, K. J., Jovanovic, T., & Stevens, J. S. (2021). Hippocampal activation during contextual fear inhibition related to resilience in the early aftermath of trauma. Behavioural Brain Research, 408, 113282. https://doi.org/10.1016/j.bbr.2021.113282

Venkatraman, A., Edlow, B. L., & Immordino-Yang, M. H. (2017). The Brainstem in Emotion: A Review. Frontiers in Neuroanatomy, 11(March), 1–12. https://doi.org/10.3389/fnana.2017.00015

Vollmer, B., Lundequist, A., Mårtensson, G., Nagy, Z., Lagercrantz, H., Smedler, A. C., & Forssberg, H. (2017). Correlation between white matter microstructure and executive functions suggests early developmental influence on long fibre tracts in preterm born adolescents. PLoS ONE, 12(6). https://doi.org/10.1371/journal.pone.0178893

Waugh, C. E., Wager, T. D., Fredrickson, B. L., Noll, D. C., & Taylor, S. F. (2008). The neural correlates of trait resilience when anticipating and recovering from threat. Social Cognitive and Affective Neuroscience, 3(4), 322–332. https://doi.org/10.1093/scan/nsn024

Wig, G. S. (2017). Segregated Systems of Human Brain Networks. In Trends in Cognitive Sciences (Vol. 21, Issue 12, pp. 981–996). Elsevier Ltd. https://doi.org/10.1016/j.tics.2017.09.006

Young, J. M., Vandewouw, M. M., Morgan, B. R., Smith, M. Lou, Sled, J. G., & Taylor, M. J. (2018). Altered white matter development in children born very preterm. Brain Structure and Function, 223(5), 2129–2141. https://doi.org/10.1007/s00429-018-1614-4

Yu, Q., Sui, J., Kiehl, K. A., Pearlson, G., & Calhoun, V. D. (2013). State-related functional integration and functional segregation brain networks in schizophrenia. Schizophrenia Research, 150(2–3), 450–458. https://doi.org/10.1016/j.schres.2013.09.016

Zalesky, A., Fornito, A., & Bullmore, E. T. (2010). Network-based statistic: Identifying differences in brain networks. NeuroImage, 53(4), 1197–1207. https://doi.org/10.1016/j.neuroimage.2010.06.041

Zwicker, J. G., Grunau, R. E., Adams, E., Chau, V., Brant, R., Poskitt, K. J., Synnes, A., & Miller, S. P. (2013). Score for neonatal acute physiology-II and neonatal pain predict Corticospinal tract development in premature newborns. Pediatric Neurology, 48(2), 123–129.e1. https://doi.org/10.1016/j.pediatrneurol.2012.10.016

